# An integrative method to unravel the host-parasite interactome: an orthology-based approach

**DOI:** 10.1101/147868

**Authors:** Yesid Cuesta-Astroz, Alberto Santos, Guilherme Oliveira, Lars J. Jensen

## Abstract

The study of molecular host–parasite interactions is essential to understand parasitic infection and adaptation within the host system. As well, prevention and treatment of infectious diseases require clear understanding of the molecular crosstalk between parasites and their hosts. As yet, experimental large–scale identification of host–parasite molecular interactions remains challenging and the use of *in silico* predictions becomes then necessary. Here, we propose a computational integrative approach to predict host–parasite protein–protein interaction (PPI) networks resulting from the infection of human by 12 different parasites. We used an orthology-based method to transfer high–confidence intra–species interactions obtained from the STRING database to the corresponding inter–species protein pairs in the host–parasite system. To reduce the number of spurious predictions, our approach uses either the parasites predicted secretome and membrane proteins or only the secretome depending on whether they are uni- or multicellular respectively. Besides, the host proteome is filtered for proteins expressed in selected cellular localizations and tissues supporting the parasites growth. We evaluated the inferred interactions by analyzing the enriched biological processes and pathways in the predicted networks and their association to known parasitic invasion and evasion mechanisms. The resulting PPI networks were compared across parasites to identify common mechanisms that may define a global pathogenic hallmark. The predicted PPI networks can be visualized and downloaded at http://orthohpi.jensenlab.org.

**Author Summary:** A protein-protein interaction (PPI) network is a collection of interactions between proteins from one or more organisms. Host–parasite PPIs are key to understanding the biology of different parasitic diseases, since predicting PPIs enable to know more about the parasite invasion, infection and persist. Our understanding of PPIs between host and parasites is still very limited, as no many systematic experimental studies have so far been performed. Efficacy of treatments for parasitic diseases is limited and in many cases parasites evolve resistance. Thus, there is an urgent need to develop novel drugs or vaccines for these neglected diseases, and thus interest in the functions and interactions of proteins associated with parasitism processes. Here we developed an *in silico* method to shed light on the interactome in twelve human parasites by combining an orthology based strategy and integrating, domain–domain interaction data, sub-cellular localization and to give a spatial context, we only took in to account those human tissues that support the parasite’s tropism. Here we show that is possible to identify relevant interactions across different parasites and their human host and that these interactions are well supported based on the biology of the parasites.

## Introduction

Parasitic diseases result in millions of deaths each year. For example, *Plasmodium falciparum* is responsible for most of the malaria deaths with an estimated 214 million cases and 438,000 deaths worldwide in 2015 [1]. Around 7 million people worldwide are infected with *Trypanosoma cruzi*, which causes Chagas disease and results in life-long morbidity and disability as well as more than 7,000 deaths per year [1]. Another highly prevalent disease, Leishmaniasis that accounts for 20 to 30 thousand deaths a year and is caused by protozoan parasites of the *Leishmania* genus [1]. Revealing the mechanisms of parasitic infection, evasion of the host’s immune system and survival are essential to identify vaccines and therapies against these neglected diseases. One last parasitic disease, which is relevant in this work is Schistosomiasis that is a neglected parasitic disease mainly caused by five species of the genus *Schistosoma*. This parasite has a prevalence estimated in 200 million people worldwide [2]. The number of treatments for schistosomiasis is limited and their efficacy as well; in many cases parasites develop resistance. Thus, there is an urgent need to develop novel drugs or vaccines.

Typically, parasites have complex life cycles with several morphological stages and infect many host cells and tissues. For that, the parasites display a resourceful capacity to live in heterogeneous environmental conditions (intra and extra–cellular parasites) and to the pressure from the hosts’ immunological response [3]. For examples, extracellular parasites remodel tissues in order to migrate and evade the immune system [4]. Likewise, intracellular parasites shape cellular processes and remodel host’s cells to adjust their niche during infection [5]. The manipulation of these processes and pathways happens through molecular interactions that the parasites use in their own advantage.

The study of molecular host–parasite interactions is essential to understand parasitic infection, local adaptation within the host and pathogenesis, in addition these complex interactions can be described as a network [6]. Pathogens affect their hosts partly by interacting with host cells proteins, which defines a molecular interplay between the parasite survival mechanisms and the host defense and metabolic systems [7]. Understanding this molecular crosstalk can provide insights into specific interactions that could be targeted to avoid the parasite’s pathogenic consequences [8]. Intra–species protein–protein interactions (PPIs) have been studied in depth and there exist large datasets containing experimentally or computationally predicted interactions [9,10]. However, the number of available datasets providing host–pathogen PPIs is limited and challenged by the intrinsic difficulties of analyzing simultaneously the host and pathogen systems in high-throughput experiments.

Thus, host–pathogen PPIs have mainly been computationally predicted using distinct strategies such as approaches based on sequence [8,11–14], structure [15,16] and gene expression [17]. Homology–based prediction is one of the most common approaches to predict host–pathogen PPIs. These approaches have been extensively used to infer intra–species interactions [10,11,18] as well as host–pathogen PPIs [13–15,19]. These methods assume that interactions between proteins in one species can be transferred to homolog proteins in another species (interologs). Using experimental PPIs and interologs were predicted 3,090 interactions between *P. falciparum* and *H. sapiens*, these interactions were grouped based on their biological, metabolic and cellular processes [15]. Homology based methods are also combined with other approaches in order to asses the quality of the interactions obtained, these additional approaches are for example random forest classifiers and interactions are subsequently filtered for parasite specific characteristics [14].

In this work, we developed an *in silico* method to unravel the host–parasite interactome across 12 human parasites, namely *Trypanosoma brucei, Trypanosoma cruzi, Trichinella spiralis, Schistosoma mansoni, Giardia lamblia, Plasmodium falciparum, Cryptosporidium hominis, Cryptosporidium parvum, Leishmania braziliensis, Leishmania mexicana, Leishmania donovani and Leishmania infantum*. Our method is based on orthology and its strengths reside in 1) incorporating high–confidence intra–species interactions, which are not limited to known physical interactions but complemented with predictions and interactions mined from the scientific literature [10], 2) using fined grained orthology assignments instead of simple sequence similarity and 3) including parasite–specific biological context such as lifestyle (uni- or multicellular) and tissue infection. The objective of integrating multiple layers of information is to reduce the number of falsely predicted interactions, increase the reliability and thereby provide a better understanding of the parasite’s molecular mechanisms. Finally, we propose that this approach can be applied to any host–pathogen system to predict relevant molecular interactions and define the context in which they unfold.

## Materials and methods

### Proteome filtering: adding parasite-specific biological context

The first step in the prediction pipeline is to filter both the host and parasite proteomes according to the specific characteristics of the studied parasite (Fig 1A). Firstly, for the interaction to happen, proteins in the parasite need to be secreted or membrane proteins depending on the type of parasite (unicellular or multicellular). When analyzing parasites such as helminths (multi–cellular and extracellular) we used the soluble secretome. Conversely, in case of unicellular parasites, we used both the soluble secretome and membrane proteins. To identify soluble and membrane proteins, we used different available bioinformatics tools to predict subcellular localization (Fig 1B). SignalP [20] was used to identify classical secretory proteins. Proteins predicted to be secreted were scanned for the presence of mitochondrial sequences by TargetP [21] and transmembrane helices by the transmembrane identification based on hidden Markov model (TMHMM) method [22]. As well, we filtered host proteins to consider only those located on the cellular membrane and extracellular space by using the COMPARTMENTS database, which provides high-confidence information on cellular localization of proteins (confidence score > 3) [23].

**Fig 1.**
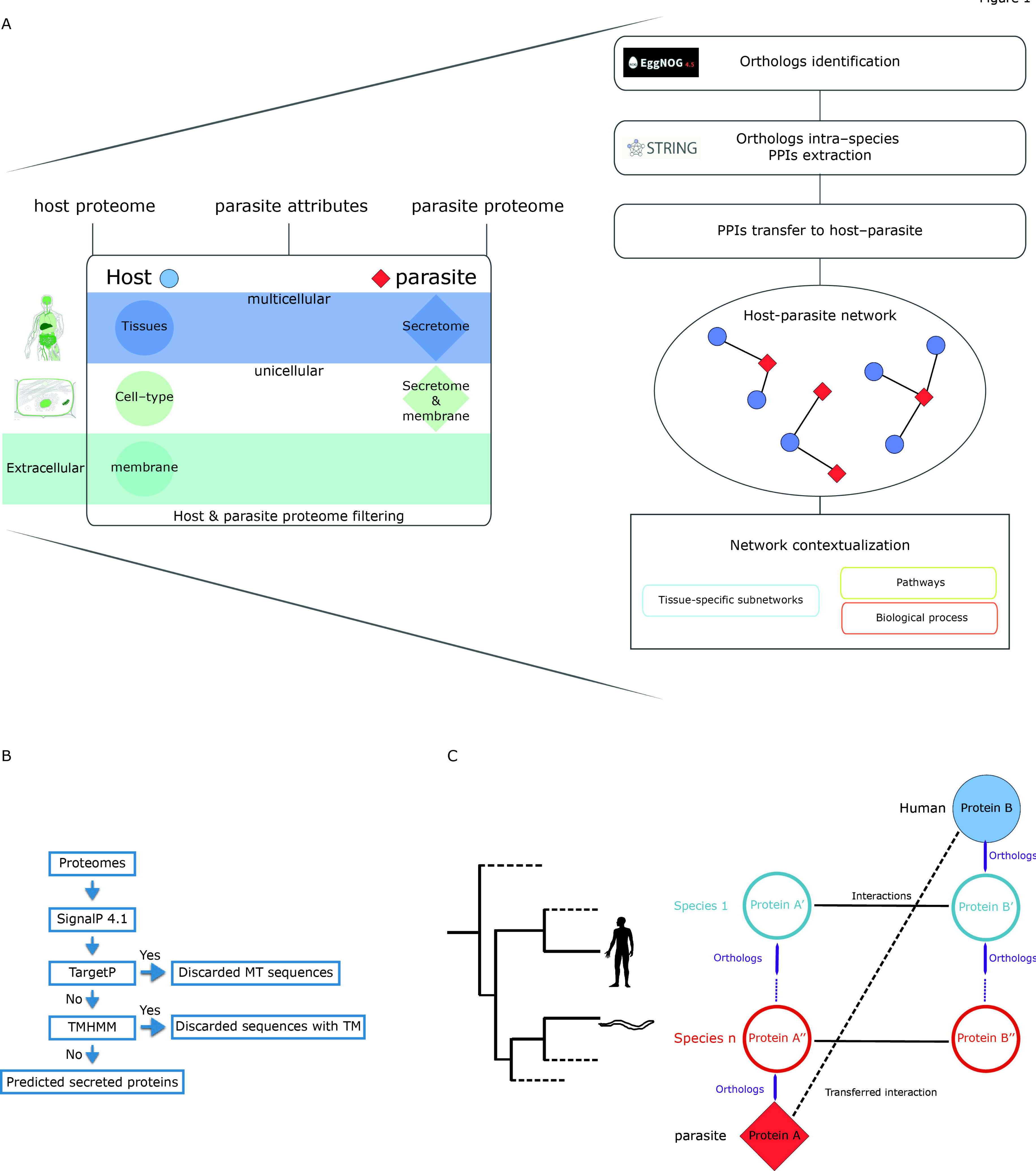
Method workflow. **A)** The method input consists of the host’s and parasite’s proteomes and specific characteristics of the parasite’s attributes (uni-multicellular) and tropism (tissues). Our approach filters the host proteome according to the specified list of tissues using TISSUES database and limits it to only cellular membrane and extracellular proteins extracted from the COMPARTMENTS database. The parasite’s proteome is filtered to only membrane or secreted proteins depending on the specified parasite attribute. The next step in the pipeline identifies in eggNOG database, orthologous proteins for both the host’s and the parasite’s filtered proteomes. The workflow continues with the extraction of intra–species protein–protein interactions from STRING database for all the orthologous proteins. These intra–species interactions are then transferred to the host–parasite system as long as the interacting proteins have an orthologous protein in host and in the parasite. **B)** Workflow for soluble secretome prediction. MT: mitochondrial, TM: transmembrane. **C)** A simple scheme of how the orthology transfer is implemented.

Additionally, our method included context on the specific tissue tropism of the studied parasite by limiting the predictions to host proteins expressed in tissues relevant in the parasite’s life cycle. For this purpose, we used the TISSUES database [24], which provides protein profiles of tissue expression. Protein–tissue associations in this database are also scored in similar way to the COMPARTMENTS database, which allowed us to use only high-confidence associations (confidence score > 3).

### Orthologs identification and PPIs transfer

Host–parasite PPIs were predicted using orthology-based transfer. This approach relies on cross-species data integration to predict inter-species protein–protein interactions. Conserved intra-species interactions from multiple organisms, namely interologs, are transferred to the host–parasite system when orthologous proteins exist in these species. For example, an intra–species interaction between protein A and protein B is transferred if the host and the parasite have orthologs to A and B (Fig 1C).

To obtain intra-species PPIs, we used the STRING database [10]. STRING provides PPIs from a variety of sources and evidence types, which expands the list of high–confidence PPIs while not limiting it to only known physical interactions. One of the integrated sources in this database is based on homology transfer, however, these transferred interactions, interologs, were omitted in our approach (STRING file including the distinction: direct *vs.* interologs). In this file, each of the evidence channels (Neighborhood, Gene fusion, Co–occurrence, Co–expression, Experiments, Databases and Text-mining) is scored independently so we combined the scores using Eq.(1).

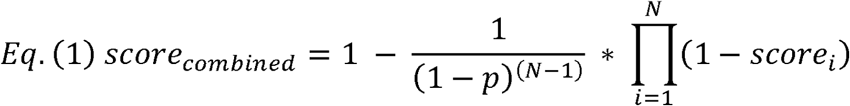

, where *i* is the different evidence channels (Neighborhood, Gene Fusion, Co–occurrence, Co–expression, Experiments, Databases and Text-mining) and *p* is the prior probability of two proteins being linked (p = 0.063). We only used high-confidence interactions (score_combined_ > 0.7).

For each of the interactions from STRING, we used the fine–grained orthologs function in eggNOG database [25] to identify orthologous proteins in human and in the different parasites. We transferred an interaction as long as it involved proteins that had an orthologous protein in the parasite and another one in human, and were in the filtered proteome. Several metrics were collected to facilitate the analyses of the interactions predicted: maximum confidence score transferred, maximum confidence score transferred from the Experiments channel in STRING, species from which the interactions are transferred and the eggNOG non-supervised orthologous groups (NOGs) they belong to, and also the number of transferred interactions corrected by the number of paralogs (duplicates).

### Domain–domain and linear motif–domain annotations

To know which interactions may be physical — rather than only functional associations — we annotated our interaction predictions with domain–domain interaction predictions from iPfam [26] and 3did [27], and linear motif–domain interactions from ELM database [28]. These databases provide predictions based on structural information from the Protein Data Bank [29]. Protein domains were predicted using Pfam scan, which combines the HMMER tool [30] and the domain models from Pfam version 29 [31]. Linear motif–domain interactions are predicted using the regular expressions provided in the ELM database.

### Network analysis

Once we obtained the predicted host–parasite PPI networks, we used the topology of the network to identify relevant proteins, which may play critical roles in the host–parasite crosstalk. We used betweenness centrality to pinpoint the proteins whose targeting would most disrupt this communication [32].

To identify key biological processes enriched in the predicted host–parasite PPI networks, we performed functional enrichment analysis using Gene Ontology biological processes (omitting the inferred evidence codes) [33] and Reactome pathways annotations [34]. We focused only on the host proteins predicted to interact given the limited functional annotation for the studied parasites. The functional enrichment was performed using Fisher’s exact test and correction for multiple testing (FDR correction Benjamini–Hochberg (BH); 0.05). The enrichment was calculated using as background only the filtered proteome.

To overcome the lack of functional characterization of the studied parasites, we also investigated the functional classification of COGs [35] provided by the eggNOG database [25]. These annotations classify COGs into functional categories that can be used to characterize the proteins grouped in these clusters (S6 Fig). These categories were transferred to the parasite proteins in the network and when the category was “Function unknown” we assigned as putative functions the categories of their interaction partners in human (S1 Table).

### Web interface

To provide access to the predictions generated with our approach, we developed a web interface for OrthoHPI (<u>http://orthohpi.jensenlab.org/</u>). This web site provides interactive, predicted host–parasite PPI networks, built with d3.js (<u>https://d3js.org/</u>), which allow the user to easily navigate the full networks as well as the tissue–specific ones. The nodes in these networks represent parasite and human proteins and their sizes correspond to their betweenness centrality in the network. The edges between nodes show predicted molecular host–parasite interactions and are weighted using the maximum score transferred from STRING database. The predictions can be downloaded in tab separated values file format (tsv) or in Graph Modeling Language format (gml), which is compatible with Cytoscape [36].

### Results and discussion

We applied our integrative orthology-based approach to predict the host–parasite PPIs networks for 12 different parasites (Table 1; Fig 1). The method returned a total of 10,096 interactions for 6,039 proteins (5,265 host proteins, 770 parasite proteins) being *T. cruzi* and *G. lamblia* the largest and the smallest predicted PPIs networks, respectively (Table 2; Fig 2A). The overview network of all 12 parasites (Fig 2A) shows that there are predicted interactions common to all parasites and parasite–specific ones. This distinction allowed us to investigate global, parasite– or genus–specific mechanisms. Our prediction method transferred most of the STRING high-confidence intra–species interactions from model organisms (*M. musculus*, *D. rerio*, *D. melanogaster*, *X. tropicalis*, etc.) (Fig 2B).

**Fig 2.**
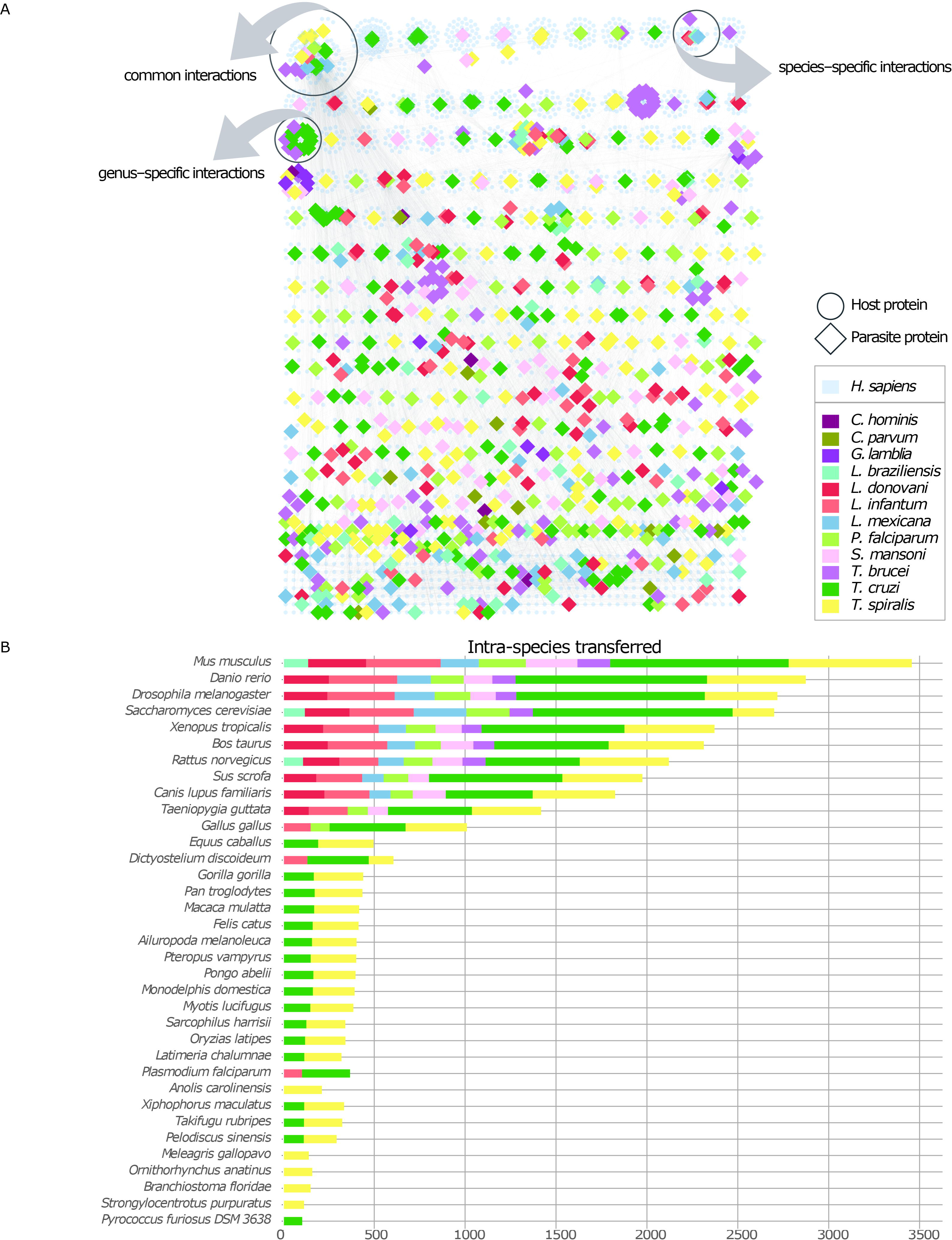
Predicted interactome. **A)** This panel shows all the predicted interactions into a single network for all the parasites and how the different parasite proteins are distributed unraveling host–parasite–specific interactions, genus–specific or shared among several parasite species. **B)** In this figure we present the species that contribute the most in the PPI transfer. Most of the interologs transferred to the host–parasite system corresponds to model organisms, which account for most of the high–confidence PPI in STRING database.

**Table 1.**
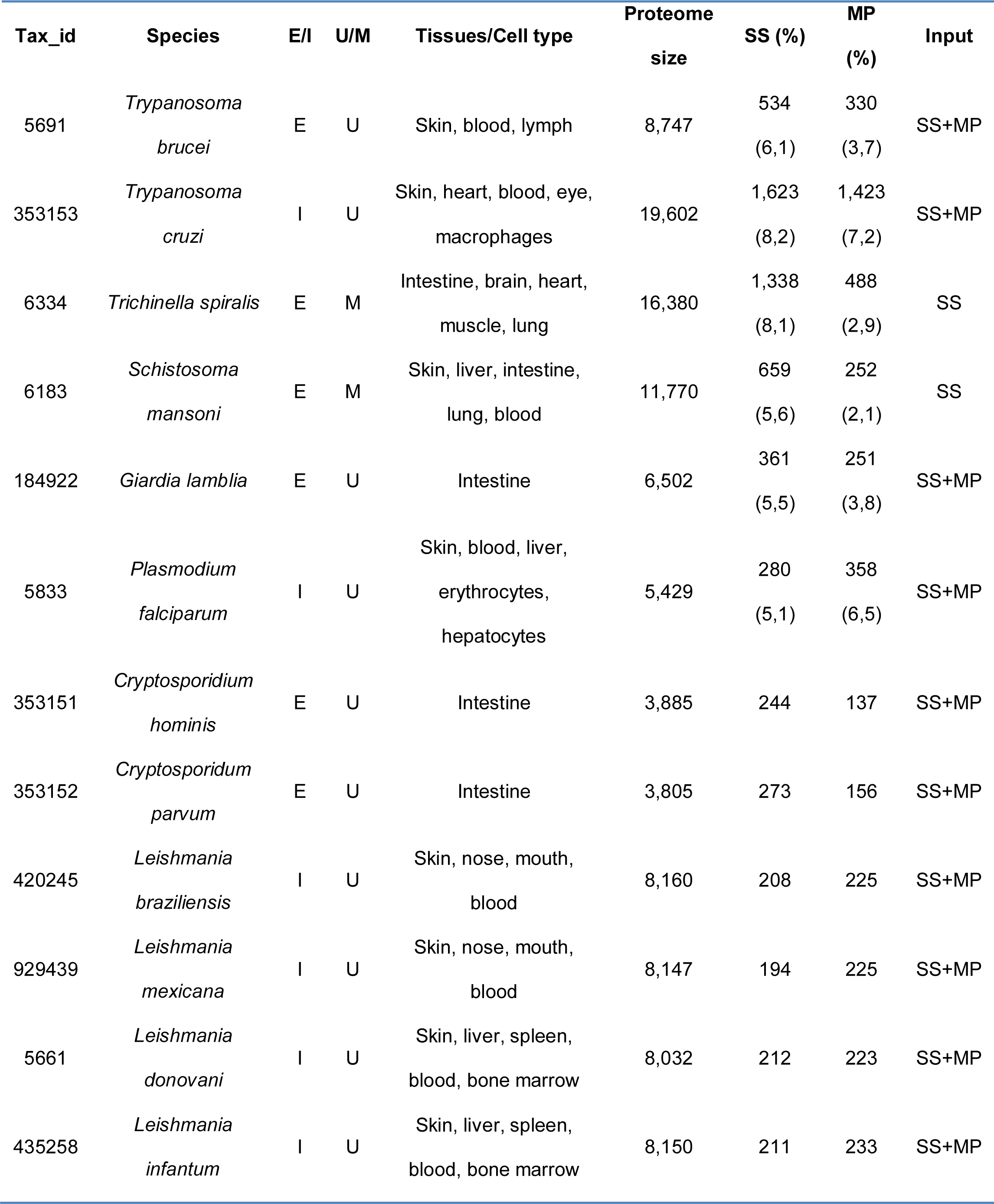
Human parasites analyzed. E: Extracellular, I: Intracellular, Cell type: host cell of an intracellular parasite. Tissues: different tissues associated with the parasite’s tropism. Proteome size: sequences in the predicted proteome. SS: Soluble secretome, MP: Membrane proteins (%: is the percentage in relation to the overall proteome.), U: unicellular, M: multicellular.

**Table 2.** Human–parasite PPI networks. This table contains the number of host and parasite nodes, the number of interactions and average degree of the predicted interactomes.

| Name | Parasite<br>proteins | Host<br>proteins | Parasite<br>nodes | Host<br>nodes | Total<br>nodes | Total<br>edges | Avg.<br>degree |
| --- | --- | --- | --- | --- | --- | --- | --- |
| *C. parvum* | 397 | 1003 | 13 | 60 | 77 | 80 | 1.039 |
| *T. cruzi* | 4056 | 6000 | 156 | 991 | 1147 | 3062 | 2.67 |
| *P.*<br>*falciparum* | 977 | 7513 | 83 | 747 | 830 | 1099 | 1.324 |
| *L.*<br>*braziliensis* | 294 | 4879 | 41 | 245 | 286 | 405 | 1.416 |
| *G. lamblia* | 658 | 1041 | 11 | 18 | 29 | 23 | 0.79 |
| *C. hominis* | 324 | 1003 | 5 | 20 | 25 | 20 | 0.8 |
| *S. mansoni* | 318 | 3503 | 69 | 429 | 498 | 581 | 1.16 |
| *L. infantum* | 325 | 8090 | 57 | 755 | 812 | 1164 | 1.433 |
| *T. brucei* | 1038 | 1824 | 101 | 265 | 366 | 528 | 1.442 |
| *L. donovani* | 310 | 8090 | 55 | 576 | 631 | 954 | 1.512 |
| *T. spiralis* | 705 | 4317 | 128 | 739 | 867 | 1389 | 1.6 |
| *L. mexicana* | 305 | 4879 | 51 | 420 | 471 | 791 | 1.677 |

The resulting host–parasite PPIs networks were analyzed to identify central proteins in the networks, tissue-specific connections and enriched biological processes and pathways. The lack of experimentally validated host–parasite interactions that could be used as a gold–standard impedes validating the quality of the predicted interactions. Instead, we validated the plausibility of the network by looking at the known functions in which the involved proteins participate. The analyses are divided into two: a case study focusing on the in-depth examination of the human–*S. mansoni* predicted interactome, and a global study of the common mechanisms targeted by the studied parasites and the shared human interactors.

### Case study: Human – *S. mansoni* interactome

#### Network topology analysis

In this analysis, we use the topological characteristics of the predicted human–*S. mansoni* PPI network to identify central proteins. Network centrality helps prioritizing proteins that have a relevant role in the network, which may translate into biologically relevant roles as well. In the human–*S. mansoni* interactome network (Fig 3) the nodes with the highest centrality were alpha-2 macroglobulin (Smp_089670), heat shock protein 70 (Smp_049550), laminin subunit beta 1 (Smp_148790) and the cell adhesion molecule Smp_171460 (Fig 4). A literature search revealed that these parasite proteins are indeed involved in crucial biological processes needed for invasion and survival of the parasite: protease activity regulation, inhibition of blood coagulation, cell adhesion and migration [37]. Our results predict that the *S. mansoni* protein Alpha-2 macroglobulin (Smp_089670) interacts with host proteins involved in matrix extracellular organization such as SERPINE1 or metalloproteases (MMP3, MMP13, MMP8 and MMP1). Interactions with the extracellular matrix (ECM) components influence several biological processes and ultimately the fate of the host cell [38]. Parasites encounter the ECM as a barrier and they deploy several mechanisms to overcome it such as cell adhesion, induction of ECM degradation and regulation of immune response [4,39,40]. For example, Alpha-2 macroglobulin is known to inhibit the predicted four proteinases by a ‘trapping’ mechanism that limits the access of the proteinases to their substrates [37]. Laminin subunit beta 1 (Smp_148790) and Smp_171460 are involved in cell adhesion and have interacting partners belonging to the metalloprotease, collagenase and laminin families.

**Fig 3.**
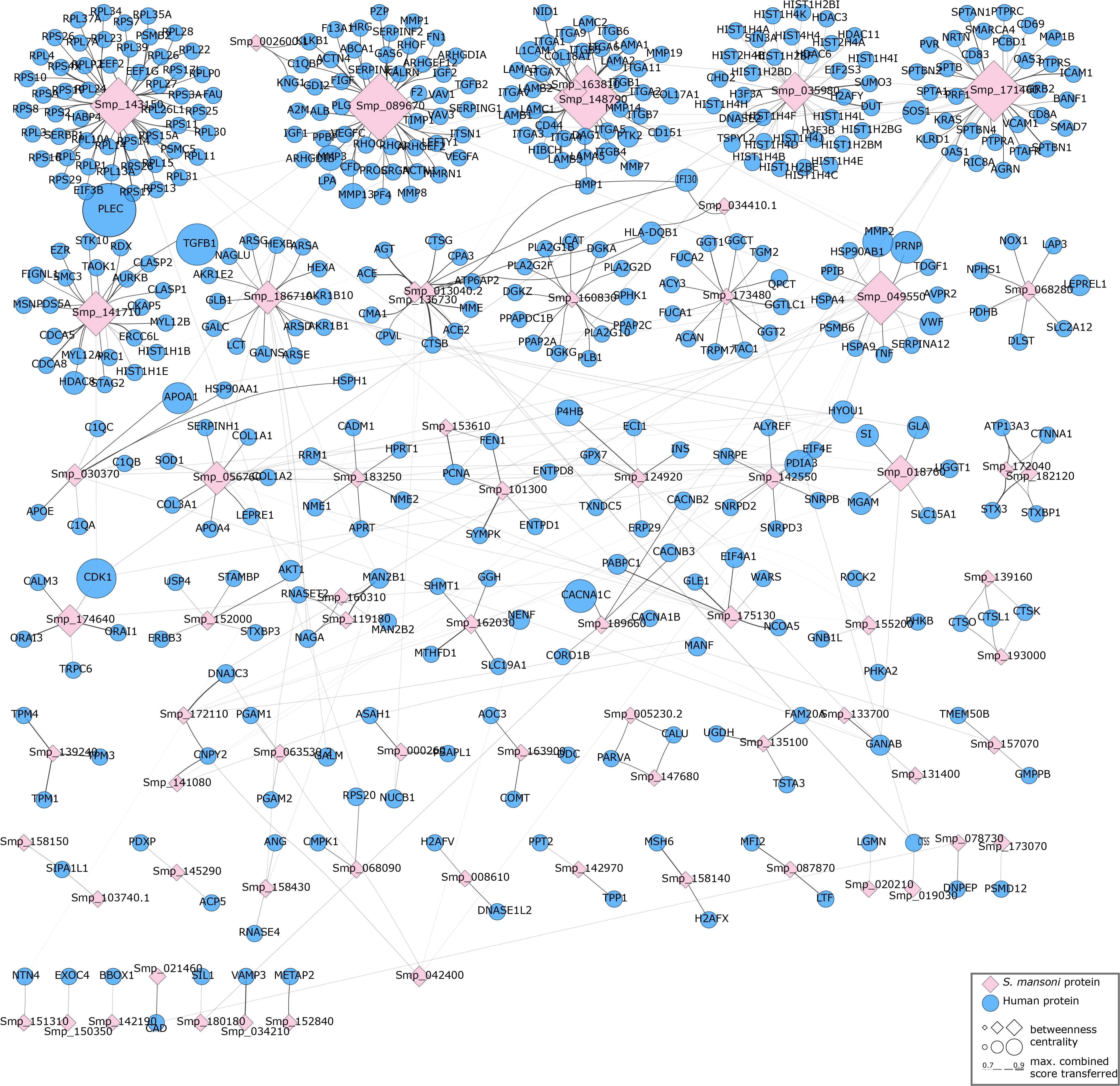
Human–*S. mansoni* predicted interactome. The layout of the nodes in the network (ellipses: human proteins, diamonds: *S. mansoni* proteins) uses Markov Clustering to group and locate the nodes. The tools used to visualize the networks in this article are: Cytoscape [36] and the clusterMaker app [41].

**Fig 4.**
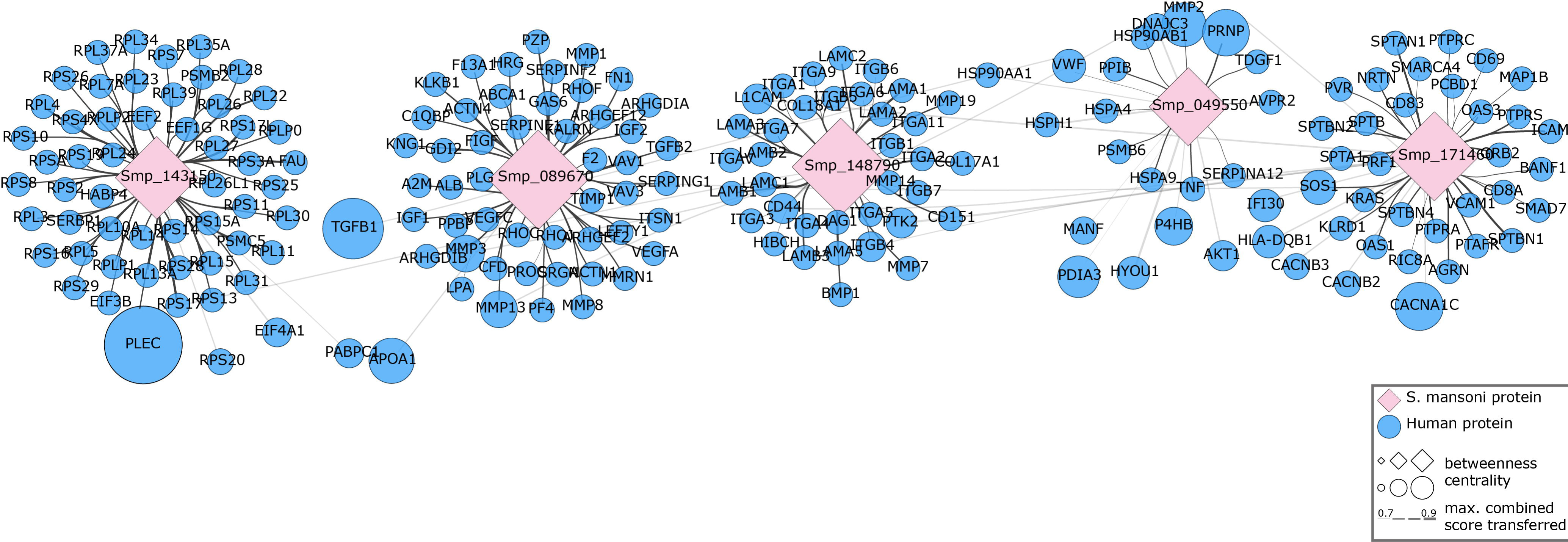
Topology analysis. This network shows the selected central nodes in the Human–*S. mansoni* predicted interactome (ellipses: human proteins, diamonds: S mansoni proteins). The central nodes are calculated using betweenness centrality and highlighted in the figure with a higher size.

The analysis highlighted as well a human protein central in the predicted interactome, PLEC protein, which strengthens cells and tissues by acting as a cross-linking element of the cytoskeleton [42]. *S. mansoni* interaction with this protein may compromise cell and tissue integrity (cell junction) and may be used to migrate across different tissues. This alteration of cell junction and tissue remodeling has been found in some helminths parasites [43].

#### Domain and linear motif analysis

We annotated the interacting proteins in the predicted human–*S. mansoni* PPI networks with domain and linear motif predictions from Pfam scan and ELM database respectively [28,30]. Around 20% of the proteins in the network (94 human proteins, 16 parasite proteins) contained interacting domains or linear motifs (Fig 5). The domain–domain or linear motif–domain interactions were integrated from iPfam, 3did and ELM databases [26–28]. The most prevalent domain in the parasite proteins is the ‘Papain family cysteine protease’ (predicted in Smp_019030, Smp_139160 and Smp_193000). This family of proteins has already been validated as drug targets for many parasites including *P. falciparum* and the African and American trypanosomes [44]. Another relevant conserved sequence is the ‘Calcineurin (PP2B)-docking motif LxvP’ present in the parasite central node Smp_171460 and the putative alkaline phosphatase Smp_171460. Calcineurin signaling manipulation provides influence over several physiological contexts such as cell adhesion, immune response, stress and post–translationally controlled events [45,46]. Similarly, several human proteins in the network contain kinase domains (AKT1, CDK1, AURKB, TAOK1, ROCK2 and STK10), which would affirm the delivered leverage of the parasite of the host cellular signaling [47–50].

**Fig 5.**
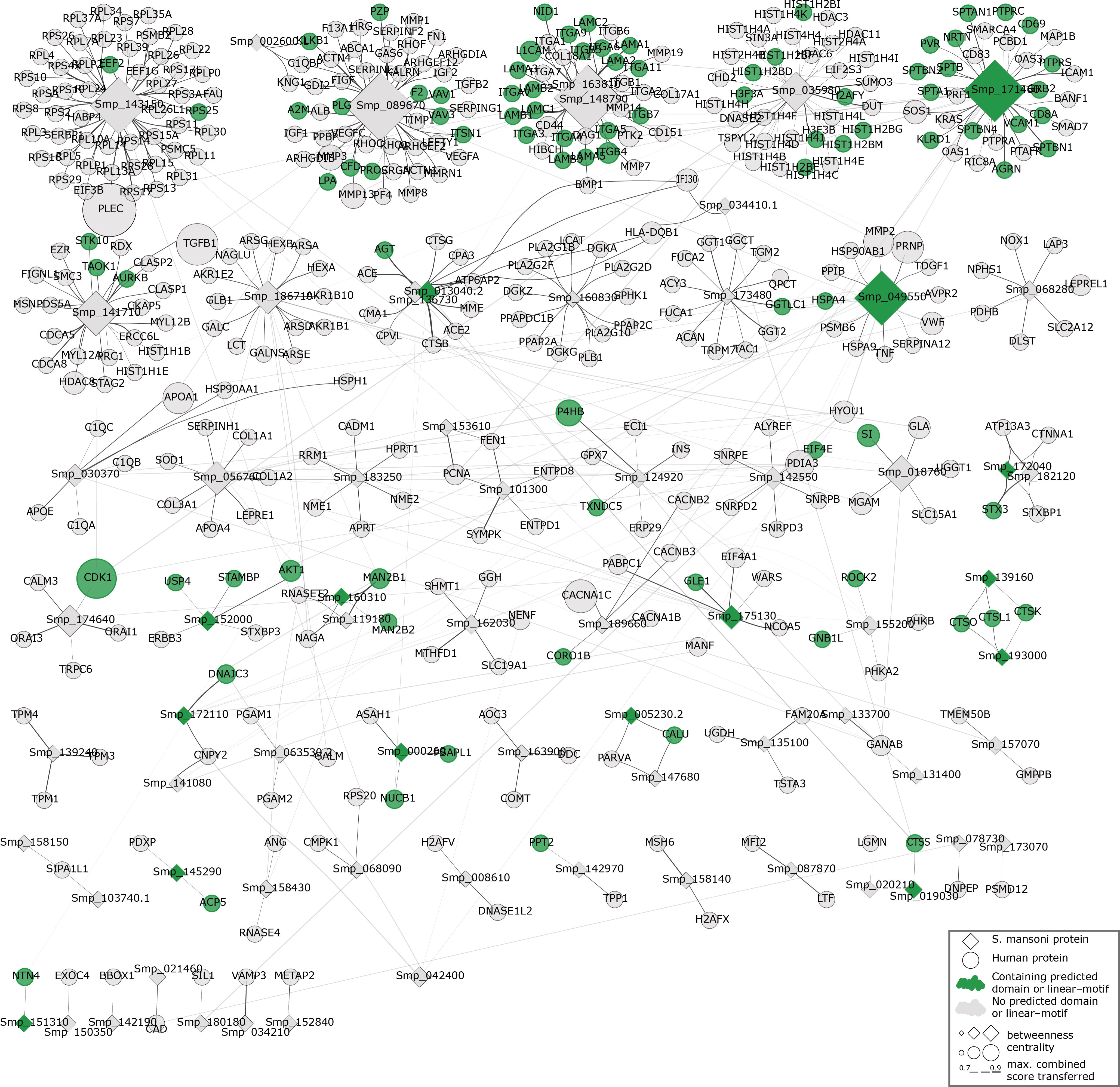
Domain and linear–motif analysis. This network shows in dark green those nodes that have a supporting domain or linear–motif predicted to interact according to iPfam, 3did or ELM. Interacting domains or linear–motifs were not found for the light green nodes. (ellipses: human proteins, diamonds: *S mansoni* proteins)

#### Tissue–specific network analysis

The human–*S. mansoni* interaction network is not static and the tissue expression data used to filter the host proteome can be used also to give context to the predicted molecular associations. Modeling spatial context into the predicted network allowed identification of relevant tissue-specific interactions through the parasite’s life cycle (S1-S5 Figs). For instance, in the blood-specific network (S1 Fig), we identified several interactions between alpha 2 macroglobulin and coagulation factors. These interactions could result in inhibition of the coagulation factors, which may facilitate the migration of the parasite through the broken tissue/vessels [37]. Equally compelling is the predicted lungs-specific interaction between the host protein FZD1, previously implicated in the inflammatory response in the alveolus [51], and the parasite protein Smp_151400 (Fig S2).

Specific characteristics of the *S. mansoni* life cycle are clearly displayed by the number of tissue-specific interactions. In this life cycle, skin is the penetration tissue and this event is happens rapidly (approximately 1 hour) [52]. According to our results, the method predicted the fewest number of interactions in this tissue. The next stage in the life cycle after skin involves migration through the bloodstream and lungs to the liver where the parasite develops into an adult male/female schistosome [53]. The number of predicted interactions in lung and liver is higher than in skin. This increase in the number of interactions could be explained by the development of the adult parasite and the energetic requirements associated with the tropism across different host tissues. The pathogenesis of human schistosomiasis initiates when the parasite’s eggs are embedded in the lung, liver or intestine, which induces inflammation, granulomas, fibrosis and subsequent morbidities [53].

### Functional analysis

#### Common pathways across parasites

The twelve parasites included in our study are strikingly divergent in terms of phylogenetic context and their biology, for example distinct forms of invasion that leads to differences in parasite’s tropism and the invasion of different sets of tissues within the host (Table 1). However, when combining all the predicted PPI networks, we found some common core pathways and biological processes targeted in the host by all the studied parasites. Indeed, it has been previously shown that evolutionary distinct parasites can target the same pathways as a result of convergent evolution, for example in *Arabidopsis* pathogens, such as the bacterium *Pseudomonas syringae* and the eukaryote *Hyaloperonospora arabidopsidis* [54]. Our analysis across the studied parasites of the targeted host pathways confirms this. We used annotations from both biological process GO terms and Reactome pathways in human to get an overview of the shared pathways targeted by the studied parasites.

In the analysis, we found terms enriched and common across 10 interactomes related to metabolism, protein folding and transport, and biosynthesis (Fig 6A). These targeted pathways have been identified already as crucial to *P. falciparum* and other Apicomplexa parasites, for instance the use of the host’s amino acid synthesis and modulation of the host metabolic network [55,56]. This host metabolism dependence has also been seen in other parasites such as in Trypanosomatids [57].

**Fig 6.**
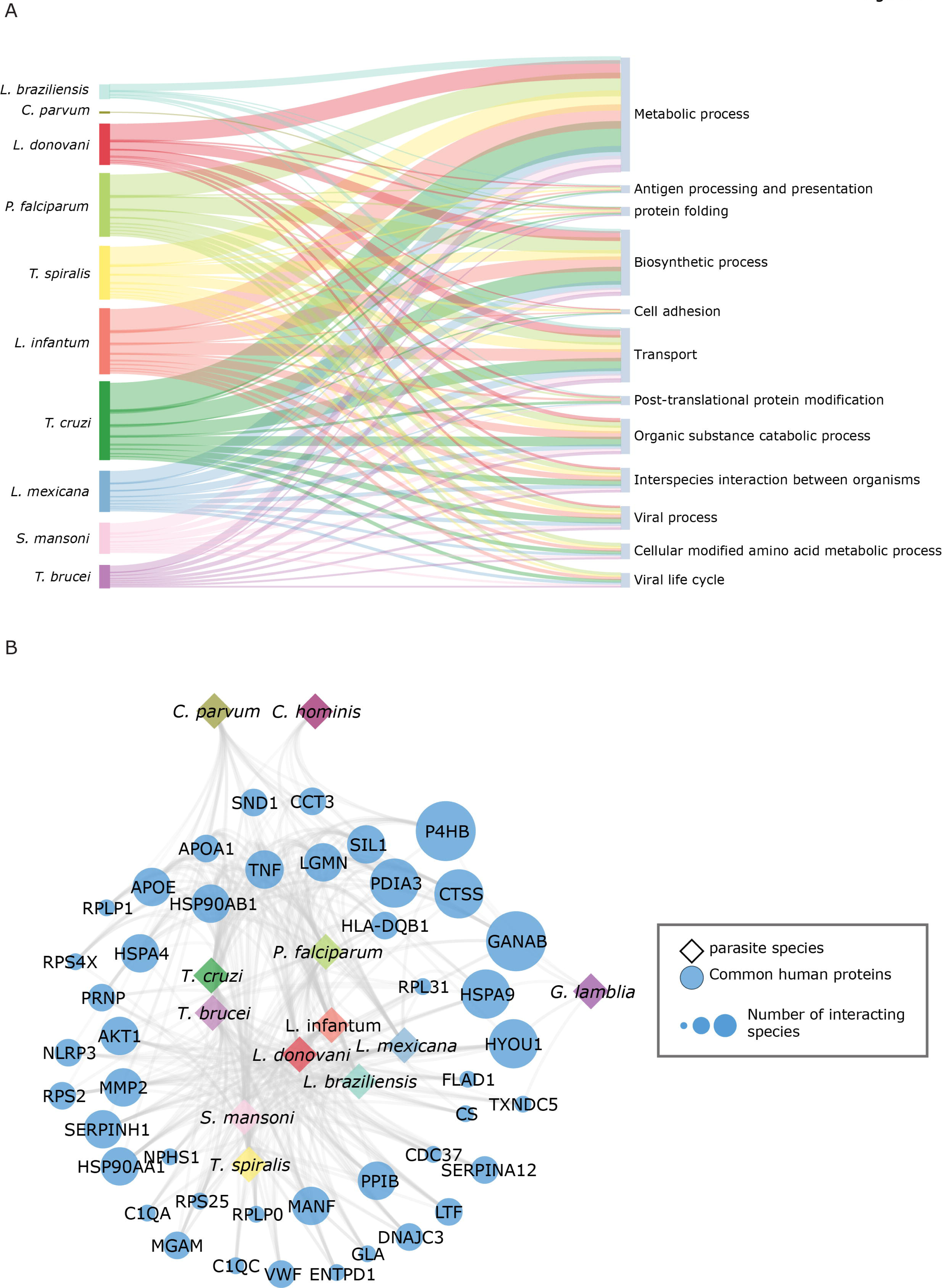
Targeted host processes and proteins. **A)** This figure shows some of the common and most relevant biological processes (GO BP) targeted by the parasites. **B)** The studied parasites interact with several common proteins some of which have been already linked to pathogenic activity. Both the targeted processes and pathways and these common host interacting proteins may open new therapeutic possibilities.

#### Pathways linked to lifestyle

Different lifestyles (intra- and extracellular parasites) may define specific targeted pathways in the host. We analyzed the common processes and pathways enriched in the intracellular and extracellular parasites specifically. The enrichment performed in the extracellular species networks did not return any lifestyle-specific targeted mechanisms, probably consequence of the common behavior of the intra– and extracellular parasites in the studied interactomes. However, the specific lifestyle requirements of the intracellular parasites (*T. cruzi*, *L. braziliensis*, *L. infantum*, *L. donovani*, *L. mexicana* and *P. falciparum*) revealed 11 exclusive biological processes related to exploitative mechanisms to acquire nutrients from the cytosol of host cells (i.e. glycerophospholipid biosynthetic process and glycerolipid metabolic process) carrying out a type of sustainability interactions. Eukaryotic pathogens are able to synthesize a number of nutrients required for growth *de novo,* however it is more advantageous to conserve energy and take host derived resources [55,58]. The enriched processes are related to the acquisition of host lipids necessary for the parasites to assemble a large amount of new membranes during replication within host cells. In terms of host immune response to parasitic infection we obtained the GO term common to intracellular parasites: antigen processing and presentation of exogenous peptide antigen via MHC class I. This term indicate that innate sensing of parasites is important for the induction of pro-inflammatory responses aimed at controlling infection [58,59].

Reactome pathways annotation provided 2 additional relevant pathways exclusive to intracellular parasites: scavenging by class F receptors and scavenger receptors (SR). These pathways include SR-A, MARCO and CD36 proteins, which belong to the innate immune defense by acting as pattern-recognition receptors, in particular against bacterial pathogens [60].

Scavenger receptors (SR) are expressed on macrophages and dendritic cells and intracellular pathogens such as hepatitis C virus or *P. falciparum* utilize the SR to gain entry into host cells [60]. The scavenger receptor MARCO is involved in *L. major* infection in macrophages [61] and the influence of other SR on *Leishmania* macrophage infection has been demonstrated [62,63]. Scavenger receptors are also involved in the formation and maturation of parasitophorous vacuole in intracellular parasites [64]. These vacuoles contain intracellular parasites initially inside the host cell and are thought to be rich in nutrients essential for parasite survival [65]. The formation of these vacuoles involves a host cell membrane invagination process, which is related to the reactome pathway Scavenging by Class F Receptors and also to the endocytic vesicle membrane [34].

#### *Leishmania*-specific pathways

Across the *Leishmania* species, we identified genus-specific biological processes related to lipid metabolic process and more specifically to fatty acid metabolism. These processes may be associated with how *Leishmania* species deplete membrane cholesterol and disrupt lipid rafts in host macrophages during the invasion process [65]. Therapies based on restoration of cholesterol levels and raft-associated proteins have been explored as promising strategies [61].

In species that cause mucocutaneous leishmaniasis (*L. braziliensis, L. mexicana*), we found 2 specific processes: regulation of acetyl-CoA biosynthetic process from pyruvate and regulation of acyl-CoA biosynthetic process. Regarding visceral leishmania (*L. donovani, L. infantum*) we found 28 specific GO terms associated with relevant biological process in the host-parasite interaction such as: regulation of defense response to virus by virus (GO:0050690), regulation of translation in response to stress (GO:0043555), glycosylceramide catabolic process (GO:0046477) among others.

In terms of Reactome pathways in *Leishmania* 4 pathways were specific. Pathways such as signal transduction (R-HSA-1227986), immune system (R-HSA-1236977), Metabolism (RHSA-1483255) to be more specific in the phosphatidylinositol metabolism, which is a component of the host cell membrane, and the other pathway was also involved in synthesis of plasma membrane intrinsic proteins (R-HSA-1660499). In mucocutaneous leishmania we obtained only 1 pathway specific, which is regulation of pyruvate dehydrogenase (PDH) complex (R-HSA-204174). Regarding visceral leishmaniasis we found 22 specific pathways related with response to stress, metabolism of proteins, disease (cancer) among others.

#### Helminth-specific pathways

In helminths (*S. mansoni, T.spiralis*) we identified 18 specific biological process, these were involved in coagulation such as hemostasis, blood coagulation and regulation of body fluid levels among others. Several helminth parasites imbibe host blood, including the hookworms, the flukes and the major livestock nematode parasites [66]. Blood coagulation is triggered by different pathways, which are targeted by blood-feeding parasites to inhibit coagulation and prolong blood flow [66]. In helminths, the inhibition of blood coagulation is related with proteinases that facilitate invasion of host tissues and digest host proteins. Additionally parasite proteinases help pathogens evade the host immune response and prevent blood coagulation [67].

Helminths-specific enriched Reactome pathways targeted host mechanisms: Sphingolipid metabolism and glycosphingolipid metabolism, which are known to have a key role in the interaction host–helminths. The sphingolipids and glycosphingolipids are a class of lipids that serve as integral components of eukaryotic cell membranes and act as signaling molecules in many cellular functions and play crucial roles in the regulation of pathobiological processes [68]. In helminths lipids may have a role in pathogenesis by helping the larvae to survive in the host tissue [68].

### Common targeted host proteins across interactomes

In this analysis, we studied the common host proteins to all or most of the studied parasites (Fig 6B), which could help identify possible general therapeutic targets. In all the inferred interactomes, GANAB (neutral alpha glucosidase AB) and P4HB (protein disulfide isomerase) proteins were predicted to interact with parasite proteins.

GANAB protein is related with the alteration of eosinophil proteome [69]. Strikingly, eosinophils are important mediators of allergies, asthma and adverse dug reactions and are also related to the host defense mechanisms against helminth infections, which are characterized by eosinophilia [70–72]. Straub et al., 2011 [69] conducted a comparative proteomic analysis of eonisophils (healthy vs hypereosinophilia from acute fascioliasis) and GANAB was one of the four proteins significantly upregulated in the *Fasciola* patient.

Protein disulfide isomerase (P4HB) has a key role in the internalization of some pathogens [73]. For example, during the host invasion process of *L. chagasi* increased levels of P4HB were found to induced phagocytosis in the promastigote phase and inhibition of expression of this gene reduced phagocytosis [74]. The role of P4HB has also been associated to other pathogens such as HIV, dengue virus or rotavirus [75–77]. In dengue virus, P4HB was linked to reduction of β1 and β3 integrins allowing for the entry of the virus [76] and in MA104 cells, thiol blockers and P4HB inhibitors decreased the entry of rotavirus [77]. These evidences show that GANAB and P4HB are relevant proteins not only in our 12 interactomes, but also in other host-pathogen systems, which confirm these proteins as possible hallmarks of the host-parasite interaction. Other host proteins common in several parasite interactomes (Fig 7) such as HYOU1, PDIA3, HSPA9, CTSS, etc., have been already found deregulated upon pathogenic infection [78–83].

**Fig 7.**
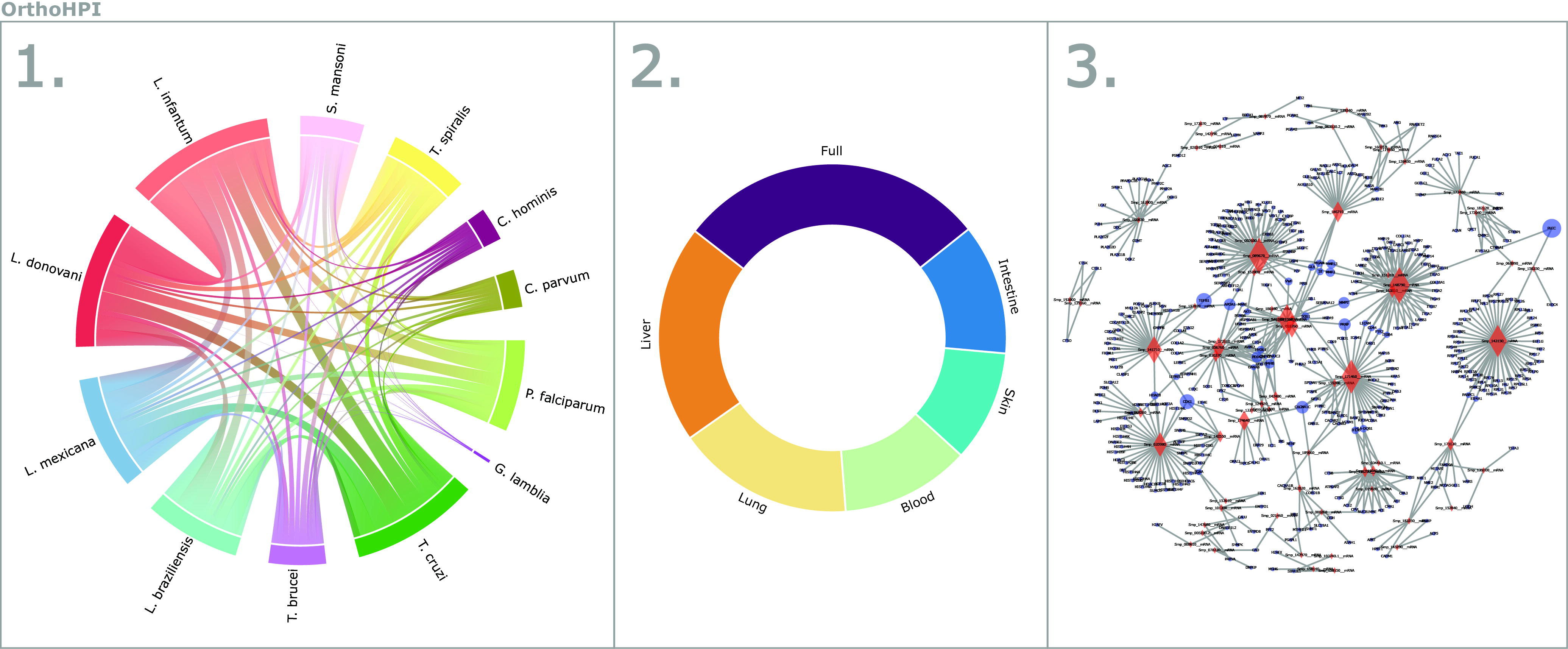
The OrthoHPI web resource. The web site (http://orthohpi.jensenlab.org) allows you to visualize the predicted interactomes for all the studied parasites in three simple steps. 1) Select parasite, 2) Choose to visualize the full network or a tissue–specific one and 3) Visualize the interactive network and download the interactions.

### The OrthoHPI web resource

We developed a web resource (<u>http://orthohpi.jensenlab.org</u>) to provide all the predicted interactomes for all the studied parasites (Fig 7). The aim of OrthoHPI is to facilitate the analysis of the predicted host–parasite PPI networks, provide easy access to the full and tissue–specific networks and produce visual, navigable and interactive networks easily manipulated by users.

## Conclusion

Our method showed molecular crosstalk between *S. mansoni* and human proteins enriched in biological processes and tissue-specific interactions essential in the life cycle of the parasite. Moreover, this approach highlighted mechanisms and specific components that may be alternative druggable targets. All these findings indicate the functional relevance of the predicted host-pathogen PPIs. An advantage of this method is that it has clear biological basis (orthology), is scalable and can be applied to many different host–pathogen systems.

Our method also could be used to assign functions to parasite’s hypothetical proteins, which are interacting with known proteins, assuming that clustered proteins tend to have similar functions and functionally related proteins can interact with each other [84–86]. This application would be useful in parasites genomics, considering that a large number of parasite’s proteins are annotated as hypothetical proteins, and our networks provides a useful resource for annotation of those proteins.

Despite the limitations that a computational method may have, our predictions provide an enriched list of host-parasites interactions, and till experimental limitations to predict host-parasite interactions are resolved, the present study or set of interactomes can efficiently serve as a theoretical base for experiments aiming to identify potential drug targets and to know more about the mechanisms to infect, migrate and persist within the host.

## Acknowledgements

This work was supported by grants to GO from the National Institutes of Health-NIH/Fogarty International Center (TW007012), GO is a CNPq fellow (307479/2016-1), and CAPES (REDE 21/2015 CAPES/MINCYT) to YCA, EMBO short-term fellowship (400-2015) to YCA. Work at The Novo Nordisk Foundation Center for Protein Research (CPR) is funded in part by a generous donation from the Novo Nordisk Foundation (Grant number NNF14CC0001).

## Supporting information

**S1 Table. Putative functional annotation of parasite proteins.** We assigned eggNOG functional annotations from host COGs to parasite proteins with no “Function unknown” in the COGs they belonged to.

**S1 Fig. Blood–specific network.** Human–*S. mansoni* predicted interactions located in blood.

**S2 Fig. Lung–specific network.** Human–*S. mansoni* predicted interactions located in the lungs.

**S3 Fig. Liver–specific network.** Human–*S. mansoni* predicted interactions located in the liver.

**S4 Fig. Skin–specific network.** Human–*S. mansoni* predicted interactions located in the lungs.

**S5 Fig. Intestine–specific network.** Human–*S. mansoni* predicted interactions located in the lungs.

**S6 Fig. Intra-species eggNOG groups functional annotation transferred.** Functions annotated in eggNOG to the most contributing clusters of orthologs groups (COGs) used in the orthology transfer method.

